# A virus-specific monocyte inflammatory phenotype is induced by SARS-CoV2 at the immune-epithelial interface

**DOI:** 10.1101/2021.09.29.462202

**Authors:** Juliette Leon, Daniel A. Michelson, Judith Olejnik, Kaitavjeet Chowdhary, Hyung Suk Oh, Adam J. Hume, Silvia Galván-Peña, Yangyang Zhu, Felicia Chen, Brinda Vijaykumar, Liang Yang, Elena Crestani, Lael M. Yonker, David M. Knipe, Elke Mühlberger, Christophe Benoist

## Abstract

Infection by SARS-CoV2 provokes a potentially fatal pneumonia with multiorgan failure, and high systemic inflammation. To gain mechanistic insight and ferret out the root of this immune dysregulation, we modeled by *in vitro* co-culture the interactions between infected epithelial cells and immunocytes. A strong response was induced in monocytes and B cells, with a SARS-CoV2-specific inflammatory gene cluster distinct from that seen in influenza-A or Ebola virus-infected co-cultures, and which reproduced deviations reported in blood or lung myeloid cells from COVID-19 patients. A substantial fraction of the effect could be reproduced after individual transfection of several SARS-CoV2 proteins (Spike and some non-structural proteins), mediated by soluble factors, but not via transcriptional induction. This response was greatly muted in monocytes from healthy children, perhaps a clue to the age-dependency of COVID-19. These results suggest that the inflammatory malfunction in COVID-19 is rooted in the earliest perturbations that SARS-CoV2 induces in epithelia.

## INTRODUCTION

Viral infections induce varied innate and inflammatory responses in the host. These responses help to control the viruses, but in some cases can become far more deleterious than the virus itself (1). Infection with SARS-CoV2 (CoV2), the cause of the current COVID-19 pandemic, leads to an upper respiratory tract infection which, if not controlled by the innate and adaptive immune responses, can evolve into a lethal pneumonia. CoV2 infection is remarkable in its clinical heterogeneity, ranging from asymptomatic to fatal (2), and several clinical characteristics demarcate the pathology associated with CoV2, when compared with other respiratory pathogens such as influenza A virus (IAV). First, critical COVID-19 is associated with multi-organ failure beyond the lungs and a concomitant severe vasculopathy (3–5). Second, bacterial coinfection, a common complication in IAV infections (6, 7), is rarely found in COVID-19, yet COVID-19 nonetheless adopts clinical aspects of bacterial sepsis (8), with an over-effusive production of inflammatory cytokines (reviewed in (9)). Finally, an important feature of COVID-19 is that children are usually spared from severe disease, showing asymptomatic or milder disease at the acute phase (10–13), even though viral loads are similar to adults (14). Such an age imbalance is not seen in IAV infections.

Many studies have aimed to understand the molecular and immunological factors that drive these clinical phenotypes (15–19). In severe COVID-19, profound alterations of the immune system have been described in myeloid cells (20, 21), along with impaired interferon (IFN) responses (22–24), impaired T cell functions (25–28), production of autoantibodies (29) and high circulating levels of inflammatory cytokines (17, 24, 30). It is not obvious how to disentangle which of these manifestations causally partake in severe pathogenesis, and which are only bystander markers of the strong inflammation. Direct pathogenicity from virus-induced damage is unlikely to be a driver, as high viral loads can exist early in asymptomatic or mild disease (31, 32), pointing to a determining role of host factors. Abnormalities in the type I IFN pathway, resulting from genetic alterations (33) or from IFN-neutralizing autoantibodies (34–37), clearly have a causative or amplifying role in COVID-19, plausibly by allowing the virus to replicate unchecked during the early phases of infection, before adaptive immune defenses can be recruited. However, the response to CoV2 involves many cellular and molecular players, and it seems likely that additional pathways beyond type I IFN underlie both resistance and pathology. More generally, the question can be framed as understanding why the newly emerging coronaviruses, including MERS and SARS-CoV1, are so pathogenic, while others that have co-evolved with humans are not. A plausible virologic explanation is that their molecular structures are mostly novel to human immune systems, as the H1N1 IAV variant was during the 1918 influenza pandemic, such that toxicity derives from immunologic novelty. Another hypothesis, not mutually exclusive, is that these highly pathogenic coronaviruses are equipped to perturb immune responses, perturbations which in turn drive severe immunopathology. Coronaviruses have large genomes, encoding many non-structural proteins, some of which are thought to have immune-modulating capabilities (38–40). They thus have the genetic leeway to evolve such strategies, their attempts at immune evasion potentially promoting particularly deleterious immunopathology.

To better understand the root factors leading to immune dysregulation in COVID-19, we designed an *in vitro* co-culture system in which immunocytes were exposed to epithelial cells infected with CoV2, then profiled by transcriptomics and flow cytometry. Epithelial CoV2 infection induced a strong, mixed inflammatory response in co-cultured monocytes resembling that of blood monocytes from COVID-19 patients. A large component of this response was not observed with two severe human pathogens used as comparators, IAV and Ebola virus (EBOV), and this response was strikingly muted in monocytes from children. Together, these results suggest that CoV2-infected epithelial cells elicit an early and specific pro-inflammatory response in monocytes, which may explain the severity of COVID-19.

## RESULTS

### In vitro SARS-CoV2 epithelial-immune co-culture induces a mixed inflammatory response in monocytes and B cells

To assess whether and how immunocytes are triggered by CoV2-infected cells, we established a co-culture model in which *ex vivo* blood immunocytes were placed in direct contact with virus-infected epithelial cells (Fig. 1A). Because primary lung epithelial cells are difficult to expand and manipulate in such conditions, we chose as a surrogate epithelial cell, the human colorectal adenocarcinoma cell line Caco-2, which is permissive for CoV2 infection (41) and DNA transfection. Under our infection conditions, CoV2 nucleocapsid (N) expression was detected in approximately 50% of Caco-2 cells by flow cytometry and immunofluorescence (Fig. 1B). Thirty-five hours after CoV2 infection of the Caco-2 monolayer, unbound virus was removed and peripheral blood mononuclear cells (PBMC) from healthy donors (HD) were added to the cultures. These were harvested 14 h later, and subpopulations were magnetically purified for transcriptome profiling by RNAseq (Fig. S1A). In good part because of the experimental requirements of BSL4 biocontainment (e.g. lysates had to be heat-treated for biosafety), RNAseq data quality was lower than customary. Rather than the usual statistical tests, identification of differentially expressed genes relied on the convergence of two independent experiments, a third experiment being used for validation (see Methods).

**Fig 1.**
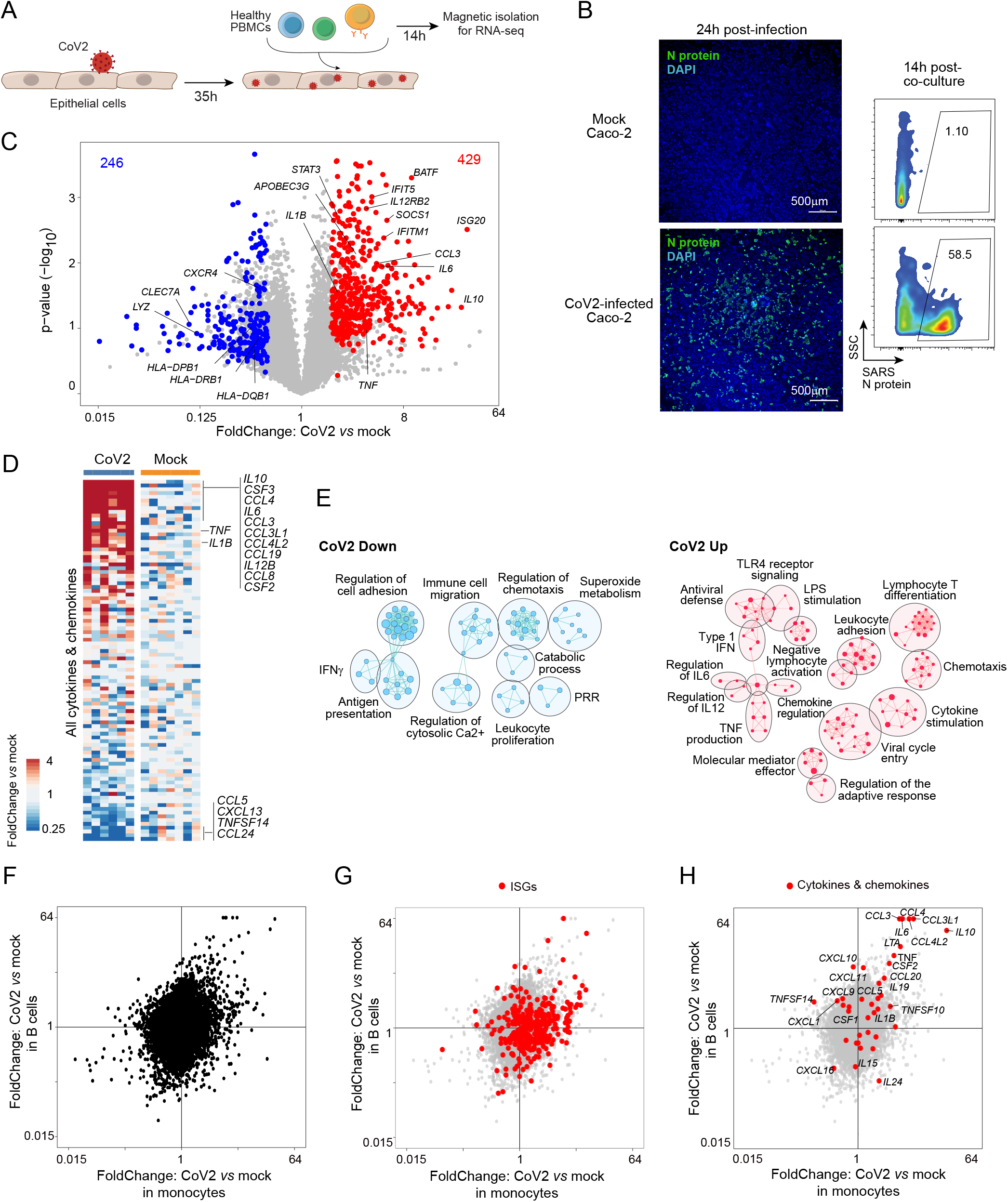
SARS-CoV2 epithelial-immune co-culture triggers a mixed inflammatory response in immunocytes. (**A**) Experimental approach. Caco-2 monolayer cultures were infected with CoV2 for two hours. Thirty-five hours post-infection, the monolayer was washed twice and PBMCs from HDs were added directly to the cultures. These were harvested 14 h later, and subpopulations (CD14+ monocytes and CD19+ B cells) were magnetically purified for population RNAseq. (**B**) Infection rate in Caco-2 cells assessed by immunofluorescence with an anti-SARS-CoV2 N antibody: displayed by microscopy at 1 day post-infection (upper panel, 4x, N antibody + anti-rabbit-AF488 + DAPI) and by flow cytometry at the time of harvesting PBMCs (∼48 h post-infection, lower panel). (**C**) FoldChange vs p-value (volcano) plot of gene expression in monocytes co-cultured with CoV2 infected Caco-2 compared to monocytes co-cultured with uninfected Caco-2 (mock). Genes from CoV2 up signature (red) and CoV2 down signature (blue) are highlighted. (**D**) Heatmap of the expression of cytokine transcripts in CoV2 versus Mock condition (as ratio to mean of mock for each condition and each experiment). (**E**) Gene ontology analyses of CoV2 up (right, red) and down (left, blue) signatures displayed as an enrichment map. Pathways are shown as circles (nodes) that are connected with lines (edges) if the pathways share many genes. Size of the node is proportional to the number of genes included in this pathway. (**F-H**) FoldChange-FoldChange plots comparing the response in monocytes (x-axis) relative to B cells (y-axis) in the context of CoV2 Caco-2 infection, without highlight **(F)** or highlighted with: interferon-stimulated genes (ISGs) (**G**); cytokine and chemokine transcripts **(H)**.

We focused the analysis on CD14^+^ monocytes and B cells, which show perturbed transcriptomes in COVID-19 patients, and are both frontline sensors of infection. In purified monocytes, a robust response was observed, with at least 675 differentially expressed genes (DEG) (Fig. 1C, which displays transcripts of the reproducible DEGs, hereafter “CoV2 signature” – Table S1). Immediately apparent were the induction of major pro-inflammatory cytokines and chemokines (*IL1B, IL6, TNF, CCL3* and *CCL4)* and a substantial number of antiviral IFN stimulated genes (ISG; e.g. *IFIT5, ISG20)*. Conversely, MHC class-II genes were significantly downregulated. Closer examination of cytokine- and chemokine-encoding genes revealed *IL10* as the most induced cytokine transcript, along with the main pro-inflammatory trio (*IL6*, *IL1B*, *TNF*; Fig. 1D). As analyzed further below, several of these traits evoked transcriptional changes in immunocytes of COVID-19 patients (15, 19). Gene ontology analysis of these genesets (Fig. 1E, Table S2) revealed a complex set of functions: cytokines, innate signaling pathways, cell mobility and adherence, and antigen presentation, suggesting that exposure to CoV2-infected cells induces profound changes in monocyte physiology. In contrast, the direct transcriptomic effect of CoV2 in Caco-2 cells was very mild (see below) with none of the changes detected in monocytes.

Analysis of B cells from the same cultures also displayed numerous changes in this setting (Fig. S1B). This response partially coincided with that of monocytes, but also included some components preferential or unique to either cell-type (Fig. 1F). Some ISGs were induced in both, although induction of the antiviral response was strongest in monocytes (Fig. 1G). Surprisingly, the cytokines and chemokines most strongly induced in monocytes were also induced in B cells (Fig. 1H). Thus, the effects of CoV2 infection on neighboring cells were apparent in several cell-types.

### SARS-CoV2 induces a stronger pro-inflammatory response compared to influenza A and Ebola viruses

Having observed a mixed inflammatory response to CoV2-infected epithelial cells in co-cultured monocytes, we next asked whether it was specific to CoV2, by comparing monocyte responses to epithelial cells infected with either CoV2, IAV (another clinically significant respiratory pathogen of the orthomyxovirus family) or EBOV (a more distant non-respiratory virus of the filovirus family, with a highly lethal hemorrhagic course also associated with strong inflammation; Fig. 2A). Epithelial infection levels were comparable between the three viruses (ranging from 30 to 80% in different IAV experiments, and ∼80% for EBOV; Fig. S2A, S2B).

**Fig 2.**
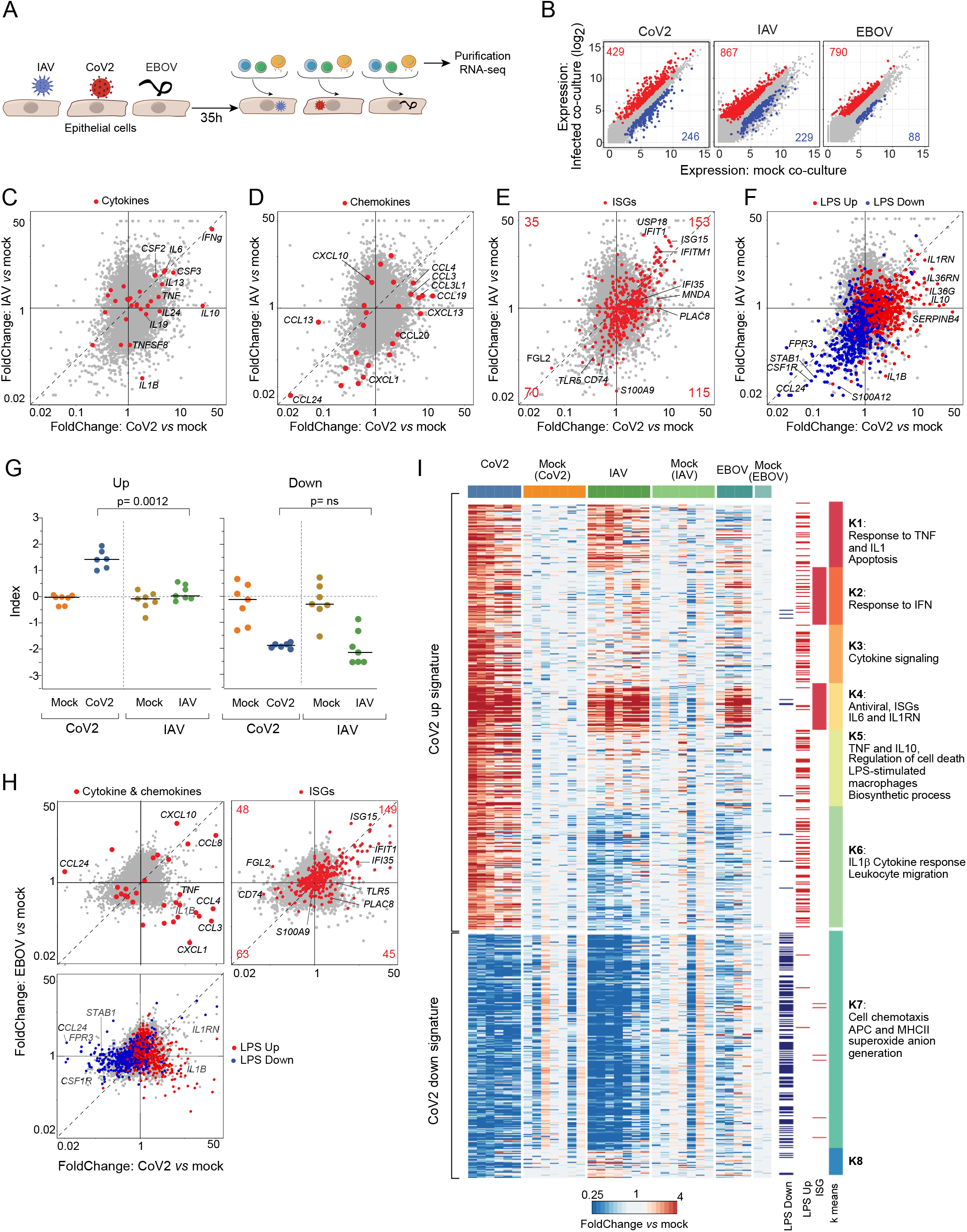
SARS-CoV2 induces a highly pro-inflammatory response compared to influenza A and Ebola viruses. (**A**) Experimental approach. Caco-2 monolayer cultures were infected with CoV2, IAV or EBOV for one, two and one hour respectively. Thirty-five hours post-infection, the monolayer was washed twice and PBMCs from HDs were added directly to the cultures. CD14^+^ monocytes and B cells were magnetically purified 14 h later for population RNAseq. (**B**) Expression-expression plots for CoV2, IAV and EBOV co-cultured monocytes vs their respective mock infections. Upregulated (red) and downregulated (blue) genes are highlighted, with numbers corresponding to the CoV2 signature as defined in Figure 1 (CoV2), or genes with a FC> 2, p< 0.01 (IAV, EBOV). **(C-F)** FoldChange-FoldChange plots comparing the monocyte response to CoV2 relative to IAV, highlighted with cytokines (**C**), chemokines **(D)**, interferon-stimulated genes (ISGs) **(E)**, and LPS upregulated and downregulated gene signatures **(F)**. (**G**) CoV2 upregulated and downregulated gene indices in CoV2, IAV, or mock infected conditions. P-values were calculated using the Mann–Whitney test. (**H**) FoldChange-FoldChange plots comparing the response to CoV2 relative to EBOV, highlighted with: cytokines and chemokines (upper left), interferon-stimulated genes (ISGs) (upper right), and the LPS upregulated/downregulated gene signatures (lower left). (**I**) Clustered heatmap of the expression of CoV2 signature transcripts, displayed as FoldChange over mean expression of mock for each condition and each batch. Each column represents one replicate. Column annotations indicate the infection condition. Row annotations indicate co-regulated modules and their dominant composition, as well as transcripts belonging to ISG and LPS gene signatures.

IAV and EBOV both induced sizeable numbers of DEGs in co-cultured monocytes (Fig. 2B), both viruses having roughly 50% stronger effects overall than CoV2). As for CoV2, the response to IAV infection in co-cultured B cells and monocytes was very similar (Fig. S2C). Direct comparison of monocyte transcriptional changes induced by CoV2 and IAV revealed that most downregulated genes were shared between the two infections, while the upregulated genes consisted of both shared and virus-specific modules (Fig. 2C-F). The CoV2-specific component included several of the pro-inflammatory cytokines described above, especially *TNF* and *IL10*; *IL1B* was even downregulated in IAV-infected co-cultures (Fig. 2C). On the other hand, *IL6* and the granulocyte/monocyte stimulating factors *CSF2* and *CSF3* were equally induced by IAV and CoV2. A set of pro-inflammatory chemokines (*CCL3*, *CCL4* and *CCL19*) were also upregulated preferentially by CoV2 infection (Fig. 2D). The eosinophil chemotactic factor *CCL24* was among the most strongly downregulated genes by both IAV and CoV2, suggesting that eosinophil recruitment is dampened in both infections. A substantial set of ISGs were induced at similar magnitudes by both viruses, but some ISGs also responded preferentially in the presence of CoV2 (Fig. 2E).

As discussed above, COVID-19 symptomatology includes several of the manifestations of sepsis, even in the absence of bacterial infection or obvious barrier breach. Furthermore, gene ontology analysis suggested that the CoV2 co-culture signature harbored elements of innate activation through Toll-like receptor (TLR) 4 activation (Fig. 1E). To test this notion, PBMCs were incubated in parallel cultures with E. coli lipo-polysaccharide (LPS), a TLR2/4 ligand. The transcriptional signature of genes induced or repressed by LPS in monocytes super-imposed strongly with CoV2-imparted changes (Fig. 2F). The LPS downregulated geneset was largely common to CoV2 and IAV infections, while the upregulated component of the LPS response was much more strongly influenced in CoV2- than in IAV-infected co-cultures (median FoldChange (FC) = 1.37 vs 0.95, chisq pval =< 0.0001 vs 0.24, respectively). Note that this intersection between the CoV2 co-culture signature and TLR activation was not merely due to dead epithelial cells released in the culture: the transcriptional changes elicited in the monocytes by exposure to lysed HEK cells (killed by freeze-thawing) bore no relation to effects of CoV2 or IAV-infected cells (Fig. S2D).

Calculating an index for responsive genes confirmed that, across monocytes from seven different HDs, the CoV2-down signature was equally elicited in CoV2 and IAV co-cultures, but that the up signature was very specific to CoV2 (Fig. 2G). To exclude that this ineffectiveness of IAV-infected Caco-2 cells to induce the full ISG set was due to suboptimal infection, we profiled monocytes in co-cultures with Caco-2 cells infected with a wide range of IAV multiplicity of infection (MOI). The CoV2-down signature was indeed most marked at an intermediate range (Fig. S2E), but the CoV2-up signature could not be significantly induced at any MOI (in contrast, ISG induction was essentially linear to infection dose).

The comparison of monocytes in EBOV- and CoV2-infected co-cultures largely reproduced the same themes (Fig. 2H): some degree of shared effects, particularly for downregulated genes (quantitatively stronger for CoV2), comparable induction of some antiviral ISGs, but a preponderance of virus-specific inductions. As in the IAV comparison, the key inflammatory cytokines and chemokines *TNF*, *IL1B, CCL3* and *IL10* were uniquely induced by CoV2 (and even repressed by EBOV). In the EBOV co-cultures, *IL6* transcripts were below the detection threshold. The LPS-induced signature showed branching into EBOV and CoV2 preferential induction, the latter being actually repressed in the EBOV co-cultures (Fig. 2H). Overall, these results are recapitulated in the heatmap of Fig. 2I, Table S4, which also highlight the dichotomy between the two ISG-containing clusters (K2 and K4), only one of which was induced in all viral co-cultures (K4), while the other is highly specific to CoV2 co-cultures (K2; Fig. 2I and Fig. S2F). Different ISGs are induced preferentially by type I IFN or IFNγ, and the ISGs of the shared cluster K4 predominantly corresponded to type I ISGs, while those of the CoV2-specific cluster K2 were enriched in IFNγ-responsive genes (Fig. S2G).

In sum, co-culture with CoV2-infected epithelial cells induces a complex response in monocytes, some of which is generic to all virus-infected cells, but most of which is quite specific to CoV2, in particular the pro-inflammatory moiety.

### Multiple SARS-CoV2 proteins can partake in triggering co-cultured monocytes

Given these specific effects of cells infected by CoV2, we then attempted to determine which viral proteins might be involved. As a screen, Caco-2 cells were transfected, in biological duplicates, with a panel of 27 plasmids encoding single viral proteins or GFP as a control (Fig. 3A). Forty-eight hour later, these transfectants were co-cultured with HD PBMCs, and the monocytes were profiled by RNAseq after 14 h. Such transient transfections can be prone to technical artefacts from cell stress during transfection, plasmid DNA, or protein over-expression (42, 43). Indeed, the RNAseq data were noisy, with substantial variation between biological replicates. We thus selected a set of differentially expressed genes whose overall variance in the dataset substantially exceeded inter-replicate variance, and with significant difference from GFP-transfected controls in at least one co-culture. We then cross-referenced these genes to transcripts of the CoV2-induced signature. Although some genes with variable expression in co-cultured monocytes showed no reproducible relation to effects in virus-infected co-cultures, two groups of genes (G2 and G6 in Fig. 3B) had very strong overlap with the CoV2-up and CoV2-down genesets. Several CoV2 proteins were able to upregulate G2 and downregulate G6 in the co-cultured monocytes (most clearly S, nsp5, 9 and 14), while others had a moderate repressive effect (N, nsp12, orf9c). No notable level of cell death was induced by any of these plasmids, and the differential effects were reproducible in parallel experiments with independent plasmid preparations (not shown). Genes in G2 mostly corresponded to the pro-inflammatory and CoV2-specific clusters K5 and K6 defined in Fig. 2. On the other hand, the ISG component of the CoV2-up signature did not belong to this group, but in a cluster with poor reproducibility and with little or no specific effects of S and nsp5 (Fig. S3A). Thus, there was a disconnect between the pro-inflammatory and the ISG moieties of the CoV2-up signature: CoV2 proteins reproduced the inflammatory but not the ISG part.

**Fig 3.**
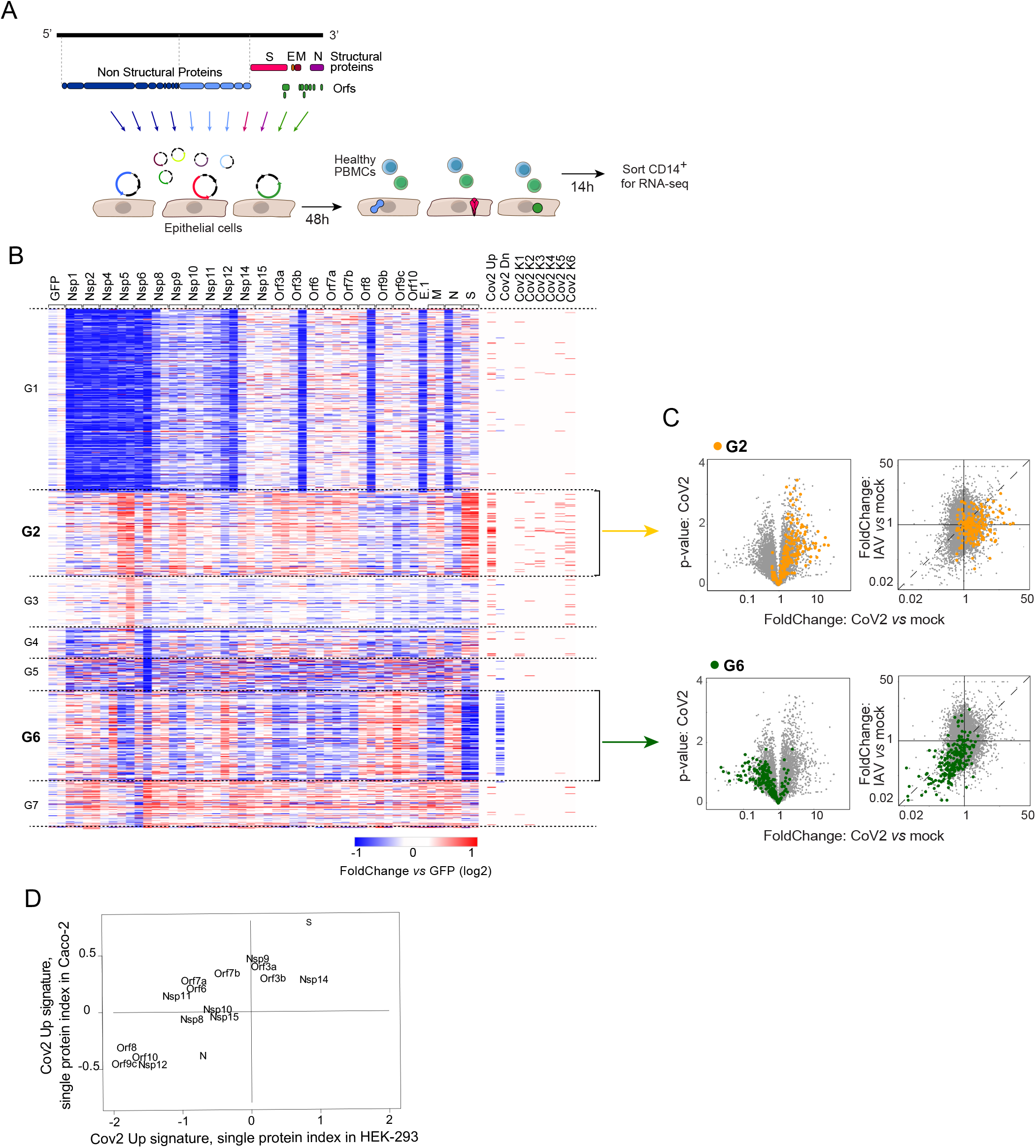
Part of the Cov2 signature can be recapitulated directly by the effects of individual CoV2 proteins. (**A**) Experimental approach. Twenty-seven CoV2 protein expression cassette plasmids or the GFP control protein expression cassette plasmid were transfected into Caco-2 or HEK cells and 48 h or 24 h later respectively, PBMCs from the same HD in each batch were added directly to the cultures. CD14^+^ monocytes were flow sorted 14 h later for population RNAseq. (**B**) Heatmap of the significantly differential expressed transcripts in monocytes co-cultured with transfected Caco-2 (selection: overall variance in the dataset substantially exceeded inter-replicate variance, and with significant difference from GFP-transfected controls in at least one co-culture FC>2 or <0.5, p<0.01). The columns represent the different transfected viral proteins (duplicates except for E). Annotations at the right ribbon show the overlap between these genes and the Cov2 signature and its different clusters. (**C**) Overlay of clusters G2 (upper panels, orange) and G6 (lower panels, green) from heatmap in panel (B) onto viral-co-culture datasets: volcano plots of gene expression in monocytes co-cultured with Cov2 infected Caco-2 compared to mock (left; as in Fig.1A) and FoldChange-FoldChange plots comparing the monocyte response to CoV2 relative to IAV (right; as in Fig.2C-F). (**D**) Scatter plot of CoV2 Up signature index in monocytes cultured with Caco-2 or HEK cells transfected with individual viral genes. Only conditions with at least 2 replicates passing QC are shown.

Reciprocally, genes from G2 and G6 identified in the transfection co-cultures proved almost entirely shifted in virus-infected co-cultures (Fig. 3C). Here again, most upregulated genes were not shared with IAV-infected co-cultures, but the downregulated signature was common to both (Fig. 3C).

For replication, we performed monocyte co-cultures with transfection into a second epithelial cell line (HEK). Comparable patterns were observed (Fig. S3B), with dominant effects of S, nsp9 and nsp14, but opposite effects of orf8,9,10, which matched genes altered in both CoV2-infected and transfected Caco-2 co-cultures (Fig. S3B, top). The effects of individual CoV2 proteins showed generally concordant distribution after transfection in both cell lines (Fig. 3D).

Thus, it was possible to replicate some of the monocyte response to CoV2-infected cells by expression of single viral proteins, confirming that the observed signatures were not merely confounders of infected co-cultures, or induced by free viral RNA. Several proteins shared the same potential, implying that changes in monocytes were not due to viral proteins acting as specific triggers, but more likely through changes that they induced in the infected epithelial cells. Active proteins settled into two groups, with diametrically opposite effects, which would presumably be balanced in the context of viral infection, but overall the virus best matched the S/nsp5/nsp14 group.

### Non-transcriptional soluble factors account for the co-culture CoV2 signature

We then attempted to tackle the mechanistic pathway though which CoV2-infected or transduced Caco-2 cells elicit the CoV2 signature in healthy monocytes. We searched for candidate mediators by examining RNAseq profiles of CoV2-infected Caco-2 cells in our cultures. Few or no genes showed significant induction, except for viral proteins themselves (Fig. 4A). In an attempt to bring out minor effects, we aligned the results of two independent culture experiments (each in biological duplicate), and observed no enrichment in the concordant segment of the graph (Fig. 4B), suggesting that most of these low-significance signals were indeed noise. The few putatively reproducible changes in Fig. 4B did not show any bias in a previously published dataset of CoV2-infected Caco-2 cells (44) (Fig. 4C). Thus, in agreement with these authors, we conclude that CoV2 infection has surprisingly minor transcriptional effects in infected Caco-2 cells. We next generated RNAseq profiles from Caco-2 cells transfected with 27 individual viral genes, and searched for transcripts that would correlate, across all the transfectants, with the ability to induce the specific signature in co-cultured monocytes. Very few transcripts showed significant correlation, with a distribution of correlation coefficient similar to that observed with random label permutation (Fig. 4D) and with no relationship between the most correlated transcripts and those putatively affected by CoV2 virus infection (Fig. 4E). We concluded that CoV2 and its proteins were inducing the activating potential in Caco-2 cells via non-transcriptional means.

**Fig 4.**
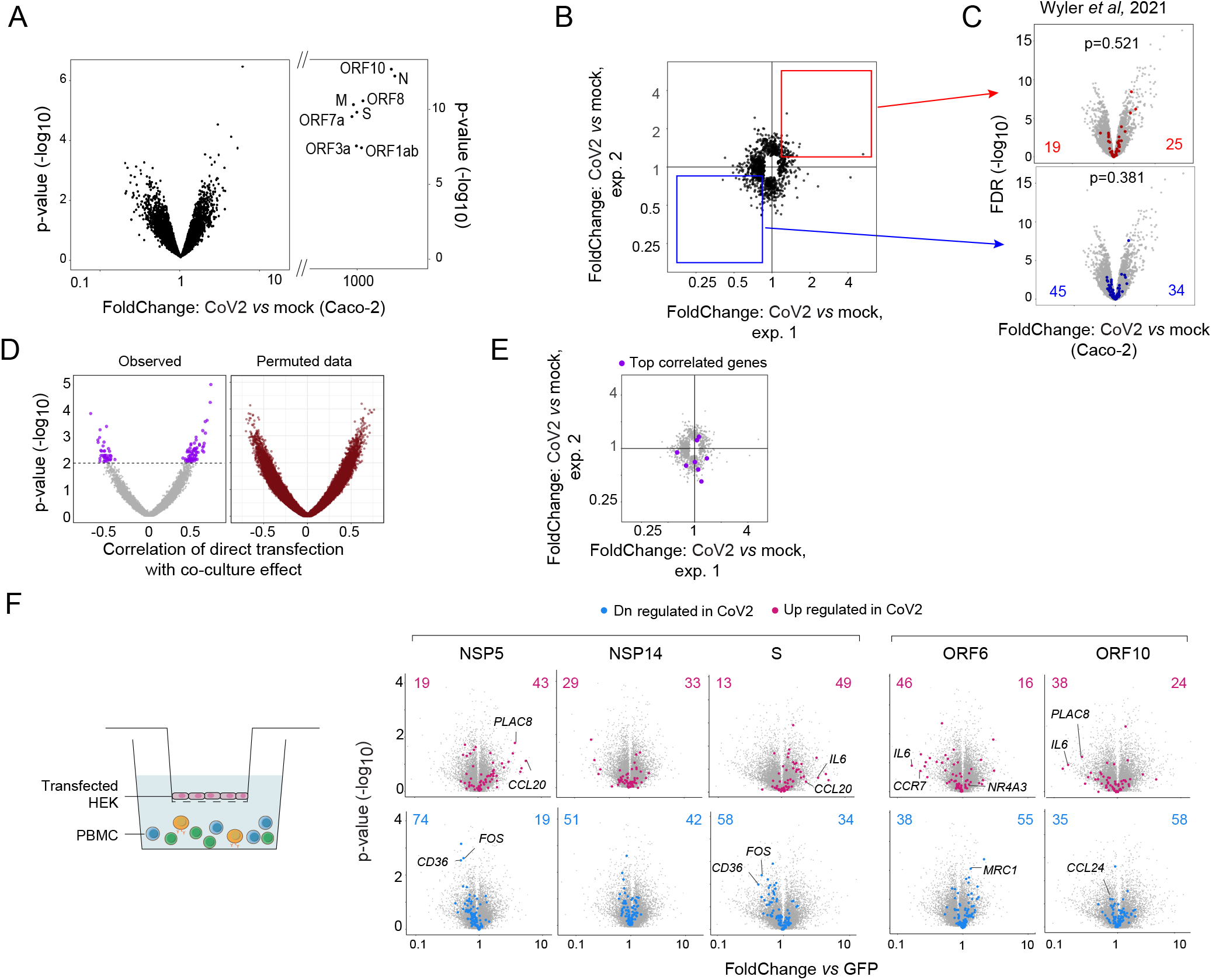
CoV2 and its proteins affect Caco2 cells non-transcriptionally. (**A**) Volcano plot of gene expression in CoV2-infected vs. mock-infected Caco-2 cells at 48h post infection. CoV2 viral genes are labeled. (**B**) FoldChange-FoldChange plots comparing the Caco-2 response to CoV2 in two independent batches (points with nominal p-value < 0.05 in either batch are shown). The red and blue squares highlight the consistent up and down-regulated changes between the two batches (|log2FC| > 0.25 in same direction). (**C**) Overlay of reproducible genes from **B** onto volcano plot of CoV2 vs mock-infected Caco-2 infected cells from Wyler et al (44). P-values of the overlaps were calculated using the Chi square test. (**D**) Per-gene correlation (Spearman) of transcriptional effect in Caco-2 cells transfected with individual viral genes with mean Cov2-Up signature index in CD14 monocytes co-cultured with transfected cells. Values obtained from data with randomized sample labels shown on right. Nominal p-values shown are based on permutation test after randomizing sample labels 100 times; purple highlighted genes: p < 0.01. (**E**) FoldChange-FoldChange plots comparing the Caco-2 response to CoV2 in two independents batches (as in **B**). The purple dots highlight the most correlated transcripts in the CoV2 protein transfected Caco-2 dataset (purple genes, p<0.01, from **D**). (**F**) Overlap of the CoV2 signature in monocytes co-cultured with transfected HEK in Transwells (HEK were cultured in the upper compartment and PBMCs were added in the lower compartment). Volcano plots of gene expression in monocytes co-cultured with transfected HEK (nsp5, nsp14, S, orf6, orf10) compared to monocytes co-cultured with GFP-HEK in a Transwell setting. Part of the Cov2 signature significantly up (red) or downregulated (blue) in HEKs is highlighted and numbers are shown.

To determine if these CoV2-related effects were mediated by cell-to-cell contact or via soluble factor(s), we used a Transwell chamber to co-culture monocytes and HEK cells transfected with a selected set of viral genes (Fig. 4F). Many of the CoV2-related effects were reproduced in this model, in particular the comparable effects of nsp5, 14 and S, implying diffusible mediators for at least some of the CoV2-provoked effects. Thus, the epithelial response to infection by CoV2 virus, or by the enforced expression of its proteins, involves the generation of soluble mediators but not ones that are induced at the transcriptional level.

### The CoV2 signature carries to monocytes of severe COVID-19 patients

These observations raised the question of the biological and clinical relevance of these *in vitro* results to the *in vivo* setting. Do these results recapitulate perturbations described in COVID-19 patients? We extracted gene expression datasets of myeloid cells from published profiling studies of COVID-19 patients and looked for reciprocal enrichment of transcriptional effects (Fig. 5A; two studies were probed in detail, but shallower examination of other studies shows the conclusions to be generally applicable). First, in myeloid cells from the lungs of COVID-19 patients (19), whose contact with infected epithelia would most closely mimic our experimental configuration, gene expression signatures of alveolar macrophages from severe patients proved upregulated in our CoV2-co-cultured monocytes (Fig. 5B, left, chisq p<10^−4^); some aligned with the swath equally affected in IAV- and CoV2-infected co-cultures (including ISGs like *IFI27, ISG15*), but the largest group belonged to the CoV2-specific quadrant (e.g. *IL1B, TNF, CCL3, CD163, TIMP1, PLAC8)*. On the other hand, genes over-expressed in macrophages from HD lung were unaffected or downregulated in our datasets (Fig. 5B, right, chisq p=0.163). In blood monocytes (15), chosen to assess a systemic spread of the effect, the index computed from the co-culture CoV2-up signature showed a clear correspondence with disease severity (Fig. 5C). In the other direction, the genes whose expression was up- or downregulated in blood monocytes from these severe COVID-19 patients relative to unexposed controls showed a strong bias in our co-culture datasets (Fig. 5D, chisq p<0.006).

**Fig 5.**
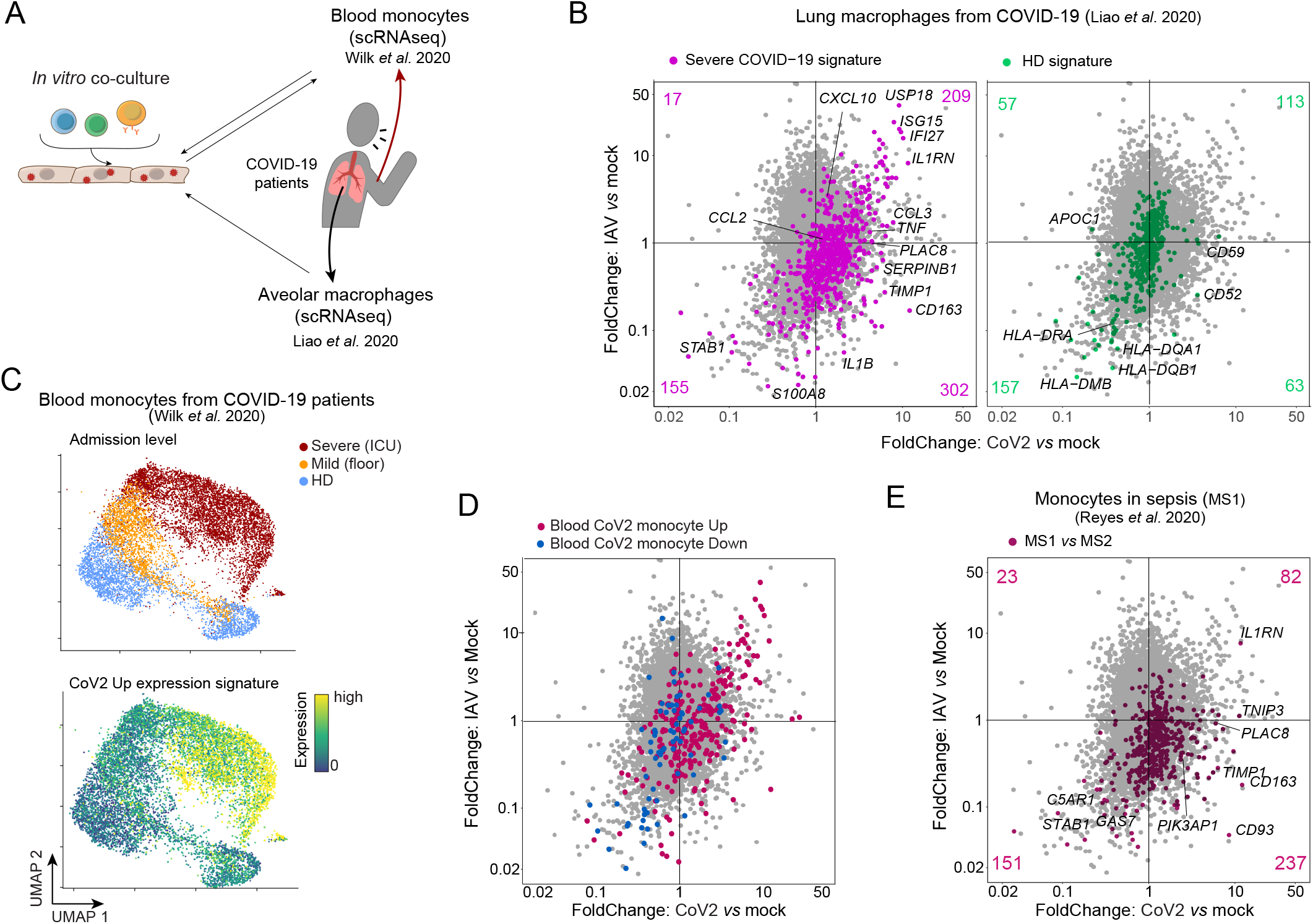
The CoV2 signature overlaps with myeloid signatures of severe COVID-19 patients. (**A**) Analysis scheme for comparison of the CoV2 monocyte co-culture *in vitro* signature with gene expression datasets of published COVID-19 studies from systemic (blood monocytes) and local (lung macrophages) myeloid compartments. (**B**) FoldChange-FoldChange plots comparing the response of monocytes co-cultured with CoV2 infected Caco-2 relative to monocytes co-cultured with IAV infected Caco-2. Highlighted gene signatures from lung macrophages from severe COVID-19 patients (left) or from HDs (right) (19). (**C**) 2D UMAP representation of CD14+ monocyte scRNAseq extracted from (GSE150728) (15). Each cell is color-coded by the patient’s severity at clinical admission (upper panel) or by the expression of the CoV2 up signature genes (lower panel). **(D-E)** FoldChange-FoldChange plots comparing the response of monocytes co-cultured with CoV2 infected Caco-2 relative to monocyte co-cultures with IAV infected Caco-2, highlighted with gene signatures from: (**D**) blood monocytes from severe COVID-19 patients (15); **(E)** MS1 (sepsis-state of monocytes) versus MS2 state (HLA-DR^hi^ monocytes) (45).

Given the described “sepsis without bacteria” clinical state of severe COVID-19 patients (3, 8) and the strong overlap between LPS-induced genes and our CoV2 co-culture signature, we asked if the CoV2 signature correlated with transcriptional alterations of the myeloid compartment in severe sepsis. Reyes et al. reported an expansion of a specific monocyte state (MS1) in patients with severe bacterial sepsis (45), which was also upregulated in monocytes from COVID-19 patients (21). Highlighting MS1 versus MS2 (classical MHC-II^high^ monocytes) signature genes into the co-culture datasets revealed a significant enrichment in the CoV2-co-cultured monocytes but a strong downregulation from co-culture with IAV-infected cells (Fig. 5E, chisq p<10^−4^). Thus, the *in vitro* co-culture CoV2 signature recapitulates the dysregulated myeloid state reported in severe COVID-19 patients, both at the local and systemic level, and overlaps with the bacterial sepsis monocyte profile.

### Monocytes from children have muted responses to SARS-CoV2

Having observed a specific response to CoV2-infected cells in co-cultured monocytes which corresponded to signatures in blood monocytes in severe COVID-19, we hypothesized that these effects might be related to age-dependent course of disease and the mostly benign evolution of COVID-19 in children. To test this notion, we co-cultured PBMCs from healthy children (4-14 years of age) with mock- or CoV2-infected Caco-2 cells, and compared monocyte transcriptional responses to those observed with adult monocytes (2 independent BSL4 experiments). Analysis of the CoV2-up and -downregulated signatures derived from adults showed that the monocyte response to CoV2 was qualitatively conserved in children (**Fig. 6A**). However, a direct comparison of adult and children monocyte responses showed a marked attenuation in children compared to adults, evidenced by the off-diagonal placement of most transcripts (Fig. 6B). This reduction applied to ISGs (Fig. 6B, right) and to key cytokine and chemokine components of the monocyte response to CoV2 (Fig. 6C, D). This shift was particularly marked for the major pro-inflammatory cytokines and chemokines induced by CoV2 in adult monocytes (*IL6, IL10, TNF, CCL3* and *CCL4*), several of which were essentially flat in children’s monocytes relative to their controls (Fig. S4). Calculating the CoV2 transcriptional index derived above confirmed that the CoV2-specific response in monocytes was significantly diminished in the monocytes of children, with no difference in responses to IAV-infected cells (Fig. 6E). Thus, in this model of initial immune encounter with infected epithelial cells, monocytes from children react in a muted fashion compared to monocytes from adults, correlating with their lower susceptibility to severe COVID-19.

**Fig 6.**
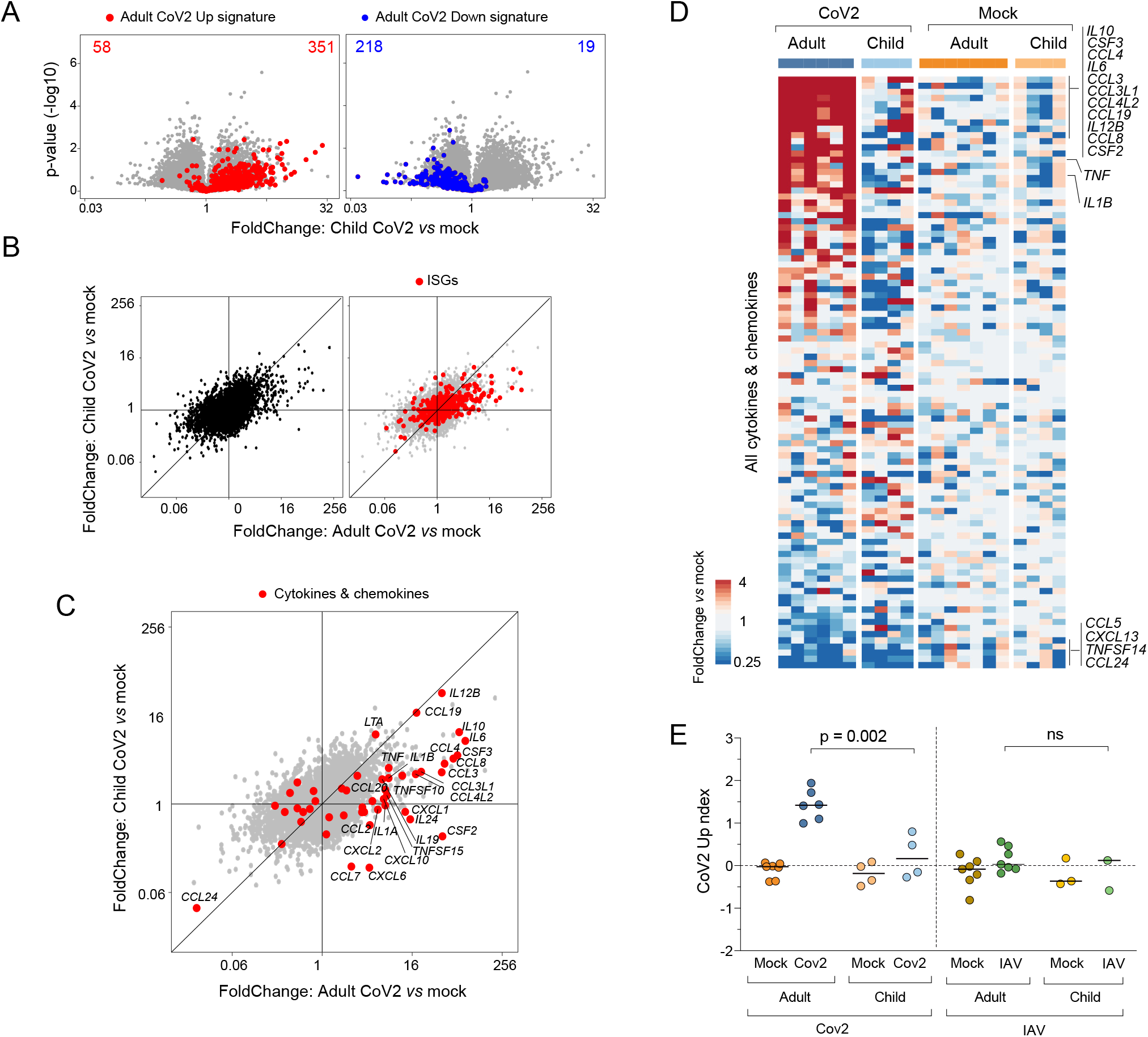
Monocytes from children have muted responses to SARS-CoV2. (**A**) Volcano plots displaying expression changes in children’s monocytes co-cultured with CoV2- or mock-infected Caco-2 cells. The adult CoV2 upregulated (red) and downregulated (blue) gene signatures are highlighted. (**B-C**) FoldChange-FoldChange plots comparing the CoV2 response of children’s vs adults’ co-cultured monocytes, without highlight (**B**, left) and highlighting the behavior of ISGs (**B**, right) and cytokine and chemokine transcripts (**C**). (**D**) Heatmap of cytokine and chemokine induction in children’s and adults’ monocytes co-cultured with CoV2-infected epithelial cells, displayed as FoldChange over mean expression of mock for each condition and each batch. Each column represents one replicate. Column annotations indicate the infection condition. (**E**) CoV2 upregulated gene index in children’s or adults’ monocytes co-cultured with CoV2-, IAV-, or mock-infected epithelial cells. P-values were calculated using the Mann–Whitney test.

## DISCUSSION

We developed an *in vitro* model of the initial encounter between immunocytes and CoV2-infected epithelial cells to investigate the cause of CoV2-induced immune changes. CoV2-infected epithelial cells directly stimulated a mixed antiviral and inflammatory response in monocytes, with components that were unique to CoV2 when compared to influenza and Ebola viruses. Several CoV2 proteins could individually recapitulate parts of this pro-inflammatory response. A comparison of monocytes from adults and children demonstrated that children have quantitatively muted responses to CoV2-infected cells.

Returning to the question of why CoV2 induces such a unique pathological response in some patients, our results offer several suggestive insights. Supporting our *in vitro* results, this unique inflammatory component has also been observed in transcriptomic analyses from severe COVID-19 patients (19, 46), whereas it seemed absent in patients with severe IAV (47). This component was shared between B cells and monocytes. The strong pro-inflammatory character of monocytes co-cultured with CoV2-infected cells, distinct from IAV and EBOV co-cultures and reminiscent in many ways of a TLR-driven response, suggests that CoV2 may deviate the immune response at its earliest stages, at the expense of effective antiviral immunity. Such an idea would nod toward a mechanistic underpinning of the “sepsis without bacteria” clinical picture of COVID-19 that has been reported (3, 8). The induction of *IL6, TNF* and *IL1B* are easy to consider in this context, but *IL10*, classically considered an anti-inflammatory cytokine, is more puzzling. Like IL6, IL10 is strongly associated with COVID-19 severity (18, 48, 49), and some prior reports suggest that it may paradoxically enhance inflammation in such settings (IL10 enhances endotoxemia (50) and induces IFNγ in patients with Crohn’s disease (51)). Later in the infection, this inappropriate initial polarization of the innate immune system may give rise to misfocused adaptive immune responses, such as the early germinal center exit of B cells and the poor T cell responses described in severe COVID-19 (18, 52, 53).

These results, and the dichotomy in ISG responses in CoV2 versus IAV/EBOV co-cultures, also speak to the role of IFN and ISGs in COVID-19 pathogenesis, some aspects of which have been difficult to integrate. Type I IFNs are clearly important in controlling CoV2 infection, as deficiencies in IFN signaling, of either genetic or immunologic origin, are strong risk factors for severe COVID-19 (33, 34, 36, 37). Accordingly, many profiling studies have reported ISG upregulation in PBMCs (for instance (15, 28, 47, 54)) or bronchoalveolar lavage (46) from severe COVID-19 patients. On the other hand, epithelial cells infected with CoV2 have been reported to produce low levels of type I IFNs (22, 55) relative other viruses, and some *in vivo* studies have observed low type I IFNs in severe COVID-19 patients (22–24). In the co-cultures, ISGs were induced along with the pro-inflammatory response, which is not surprising given that the response to CoV2-infected cells is highly reminiscent of the response to TLR ligands, and that IFN induction is a consequence of activation by many TLRs (via TRIF and IRF3). It was interesting that only half of the ISGs induced by CoV2-infected cells were shared with EBOV or IAV infection. This cluster K4 included many key anti-viral ISGs, and we propose that these correspond to true IFN-induced responses elicited by all viruses. On the other hand, the CoV2-specific ISGs of cluster K2 may be induced independently of Type-1 IFN, e.g. by IFNγ or through other signaling pathways directly activated by infected epithelial cells. Thus, from the initial interaction between CoV2-infected epithelial cells and monocytes, the stage is set to counterbalance an IFN response that is essential for viral clearance by a pro-inflammatory diversion.

How do CoV2-infected cells stimulate this CoV2-specific response in monocytes, and what is the molecular mediator(s)? Our screen of individual CoV2 proteins and comparisons to LPS offer some clues: (i) The strong overlap with responses induced by LPS (a TLR2/4 ligand) implies that signaling pathways downstream of TLRs are being triggered in monocytes and B cells (ii) It is likely not the viral proteins themselves that activate these pathways, since several CoV2 proteins (as different as the ACE2-binding Spike protein and the viral protease nsp5) have superimposable abilities, this in Caco-2 as well as HEK cells, making it difficult to envisage this explanation. In addition, there is an interesting symmetry, where proteins like Orf8 or Orf9 actually repress S-induced genes (and induce S-repressed ones), suggesting two cellular states whose balance is perturbed by viral elements. (iii) Our experimental set-up should have avoided direct CoV2 infection of the monocytes themselves, and indeed few reads from the CoV2 genome were observed in the monocyte RNAseq (unlike IAV co-cultures, where high viral reads suggested some degree of re-infection, Table S3). (iv) Soluble mediators are at least partially involved, since co-cultures with physical separation of the cells in Transwell reproduced these effects, but not simply by transcriptional induction of cytokine or chemokines, as evidenced by extensive profiling of the infected or transfected Caco-2 cells themselves. Integrating these threads, we suggest that CoV2 infection, and/or the expression of individual CoV2 proteins, causes the epithelial cells to display or release increased amounts of mediators that activate innate sensors in monocytes. Candidates include mediators such as HMGB1 (56), F-actin (57), or other cell-derived “damage-associated molecular patterns” (DAMPs (58)). Our hypothesis that host products from infected cells trigger monocytes complements recent reports of a molecular interaction between CoV2 proteins and TLR2 or C-type lectins on myeloid cells (59, 60); some of us have also reported that the S protein from SARS-CoV1, expressed in PBMCs via a herpes viral vector, can induce IL6 expression (40). This direct triggering by S may parallel the more general pro-inflammatory pathway induced by a variety of viral proteins, underlining the evolutionary importance of this response for highly pathogenic coronaviruses. Finally, Reyes et al. have shown that IL6 alone can induce in monocytes transcriptional changes (the “MS1” program) with similarities to deviations observed in COVID-19 or sepsis patients (21), suggesting a causal role for IL6. This cannot be the case here (no IL6 was detected in infected or transfected Caco-2 cells), but it may be that IL6 acts as a feed-forward loop, induced by CoV2-infected cells and then further amplifying the deviation.

Finally, what to make of the muted response to CoV2-infected cells in monocytes from children, affecting both the ISG and pro-inflammatory components? The relative protection children enjoy from severe COVID-19 is one of the most unique aspects of CoV2 compared to other common respiratory viruses (11–13). Although this is only a 2-point correlation, we speculate that the low responsiveness of their monocytes could be a key element of children’s relative protection. Mechanistically, immunocytes from children may be less responsive due to a relative naiveté vis-à-vis prior inflammatory exposures (a relative absence of “trained immunity”), or the difference may reflect the systemic pro-inflammatory tone that develops with aging (61, 62).

In conclusion, these results indicate that the dangerous inflammatory course followed by COVID-19 may be rooted in the very first immune interactions, with amplifying deviations that children are able to avoid. Modulating this inflammatory seed might prevent the subsequent exuberant and deleterious immune activation.

## MATERIALS AND METHODS

### Viruses

SARS-CoV2 stocks (isolate USA_WA1/2020), Influenza A PR8-GFP virus (A/Puerto Rico/8/1934(H1N1) and EBOV (isolate Mayinga) were grown in Vero E6 cells and purified by sucrose ultracentrifugation and titers determined. Work with EBOV and CoV2 was performed in the NEIDL BSL-4 facility. Caco-2 cells were seeded and infected 24 hrs later with CoV2 or EBOV at a nominal MOI of 10, with IAV at nominal MOI ranging from 0.1-10. After an adsorption period (2h for CoV2 and EBOV, 1h for IAV), the inoculum was removed, replaced with fresh media, and cells incubated for 35 h prior to coculturing with PBMC.

### Transfections

CoV2 expression plasmids were kindly provided by D. Gordon and N. Krogan (UCSF, San Francisco) (38). Two independent preparations of plasmids were used in independent transfection experiments. 24 hours after seeding, epithelial cells were transfected using Lipofectamine™ 3000 (Thermo Fisher), washed after 8-12h, and grown for another 24h (HEK) or 48h (Caco-2) before addition of PBMCs or lysis for RNAseq.

### Cocultures

PBMCs were collected from 17 healthy adults (21-65 years old) and 11 children (aged 4-14), either before December 2019 or without recent COVID-19 symptoms plus negative PCR within 3 days prior. These experiments were performed under IRB protocols IRB-P00021163, MBG2020P000955 and IRB15-0504. 35h post-infection or 24-48h post-transfection, frozen PBMCs were thawed and washed, then added to washed epithelial cells for 14h of coculture. For Transwell, HEK cells were transfected and re-plated onto Transwell inserts. Media from both chambers was replaced 24 hrs later, and PBMCs added to the bottom chamber for 14h co-culture. CoV2 and IAV co-cultures were performed in three independent experiments (one pilot, one main experiment and one replication experiment) with at least 3 biological replicates per condition (PBMCs from different donors). EBOV co-culture was performed for one experiment with 3 PBMC sources. Transfectant co-cultures were performed in two independent experiments, each including 2 biological replicates. Co-cultured monocytes were purified by magnetic selection (if with virally-infected cells) or by flow cytometry (if with transfected cells) prior to **l**ow-input RNA-seq per ImmGen protocol (www.immgen.org); Viral reads were mapped to CoV2, EBOV and IAV sequences from NCBI.

The data reported in this paper have been deposited in the Gene Expression Omnibus (GEO) database under accession no. GSEXXX (human RNAseq). Further details are available in *Materials and Methods*.

## ACKNOWLEDGMENTS

We thank Drs. D. Gordon, N. Krogan, T. Chatila, Y. Kai, B. Chen, J. Kagan, R. Medzhitov, V. Dixit, M. Reyes, N. Hacohen, C. Blish and M. Merad for insightful discussions, data or materials, and the Broad Genomics Platform for RNAseq during difficult times. We thank D. Ischiu for flow sorting, E. Suder and M. White for technical assistance; H. Feldmann for providing Ebola virus, and CDC’s Principal Investigator N. Thornburg and the World Reference Center for Emerging Viruses and Arboviruses for providing SARS-CoV2. This work was funded by grants from: the Massachusetts Consortium for Pathogen Readiness (MassCPR) to CB, DMK and EM; P01 AI098681 to DMK; R21 AI135912 and R01 AI133486 to EM; R24 AI072073 to the ImmGen consortium. SG-P was supported by a fellowship from the European Molecular Biology Organisation (ALTF 547-2019), JL by an INSERM Poste d’Accueil and an Arthur Sachs scholarship, KC and DAM by NIGMS-T32GM007753 and a Harvard Stem Cell Institute MD/PhD Training Fellowship.

## SUPPLEMENTARY MATERIALS AND METHODS

### Peripheral blood mononuclear cells samples

PMBC samples from 17 healthy adults (21 to 65 years-old) and 11 children (4 to 14) were all collected either before December 2019, or with no recent symptoms consistent with COVID-19 and a negative PCR test within 3 days prior to collection. Some samples were purchased frozen from AllCells (Alameda,CA,USA), or were residual materials from prior studies at Boston Children’s Hospital (EC), Mass General Hospital (LY) or Harvard Medical School (CB). These experiments were performed under IRB protocols IRB-P00021163, MBG2020P000955 and IRB15-0504.

### Cells and Cell culture

HEK (ATCC CRL-1573) and Caco-2 cells (ATCC HTB-37) were cultured in Dulbecco’s modified Eagle’s medium (DMEM, Gibco) with 10% FBS, penicillin (50 U/ml), streptomycin (50 mg/ml), and 1% MEM non-essential amino acids in a humidified incubator at 37°C with 5% CO2. Cells were passaged using 0.05% Trypsin-EDTA solution. Cultures were verified to be free of mycoplasma contamination using MycoAlert Mycoplasma Detection Kit (Lonza).

### CoV2 and EBOV propagation and titration

SARS-CoV2 stocks (isolate USA_WA1/2020, kindly provided by CDC’s Principal Investigator Natalie Thornburg and the World Reference Center for Emerging Viruses and Arboviruses (WRCEVA)) and EBOV (isolate Mayinga, kindly provided by Heinz Feldmann, NIH NIAID Rocky Mountain laboratories) were grown in Vero E6 cells (ATCC CRL-1586) and cultured in DMEM supplemented with 2% FBS, penicillin (50 U/ml), and streptomycin (50 mg/ml). To remove confounding cytokines and other factors, viral stocks were purified by ultracentrifugation through a 20% sucrose cushion at 80,000 x g for 2 h at 4°C (1). Viral titers were determined in Vero E6 cells by tissue culture infectious dose 50 (TCID_50_) assay and calculated using the Spearman-Kärber algorithm.

All work with EBOV and CoV2 was performed in the biosafety level 4 (BSL-4) facility of the National Emerging Infectious Diseases Laboratories at Boston University, Boston, MA following approved SOPs and inactivation procedures.

### Influenza A virus propagation and titration

Influenza A PR8-GFP virus (A/Puerto Rico/8/1934(H1N1), hereafter IAV) and Madin-Darby canine kidney (MDCK) cells were provided by D. Lingwood (Ragon Institute). IAV stock was made using MDCK cells and titer was determined by plaque assay as described previously (2).

### Viral infection of Caco-2 cells for co-culture

One day prior to infection, Caco-2 cells were seeded at a density of 10^5^ cells per well of a 24-well tissue culture plate or 25.10^3^ cells per well of a 96-well tissue-culture plate. Twenty-four hours later, cells were infected with CoV2 or EBOV at a nominal MOI of 10, with IAV at nominal MOI ranging from 0.1-10. After an adsorption period (2h for CoV2 and EBOV, 1h for IAV), the inoculum was removed, replaced with fresh media (2% FBS supplemented DMEM), and cells were incubated at 37°C for 35 h prior to coculturing with PBMC.

### Plasmids and transfections

Expression plasmids were kindly provided by D. Gordon and N. Krogan (UCSF, San Francisco): 27 plasmids encoding CoV2 single viral proteins and GFP as a control (3). Most proteins were cloned in a pLVX-Puro backbone under an EF1a promoter; Spike was cloned in a pTwist backbone under an EF1a promoter. They were transformed into One Shot™ Stbl3™ competent *E. Coli* (Thermo Fisher) and were amplified using Zippy plasmid miniprep kits (Zymo Research). Two independent preparations of plasmids were made and used in independent transfection experiments. Epithelial cells, in suspension (Caco-2) or seeded 24 h prior (HEK), at a concentration of 25.10^3^ cells/well, were transfected with 100 ng of each plasmid using Lipofectamine™ 3000 reagent (Thermo Fisher) in Opti-MEM™ medium (Gibco) and washed 8 – 12 h later. They were then grown for 24 h (HEK) or 48 h (Caco-2) in 96-well flat-bottom tissue culture plates before addition of PBMCs or direct lysis for RNAseq

### Coculture of PBMCs with infected or transfected epithelial cells

The same experimental design was used for viral-co-cultures and transfectant-co-cultures. 35 h post-infection or 24-48 h post-transfection, adherent epithelial cells were washed gently twice with standard culture media (DMEM, 10% FBS, penicillin (50 U/ml), streptomycin (50 mg/ml), and 1% MEM non-essential amino acids) to remove cell debris. Frozen PBMCs were thawed in a 37°C water bath for 90-120 sec then added dropwise to 9 mL of pre-warmed culture medium and centrifuged at 300 x g for 7 min. After removing supernatant, the PBMC pellet was resuspended in pre-warmed culture medium at a concentration of 1.5.10^6^ cells/ml. This PBMC suspension was slowly added to the epithelial cells at a final concentration of 7.5.10^5^ PBMCs/well (24-well plate, infections) or 2.10^5^ PBMCs/well (96-well plate, transfections). PBMCs and epithelial cells were then co-cultured for 14 h in a 37°C incubator (5% CO2).

For Transwell transfectant experiments, HEK cells were transfected in standard flat-bottom plates, dissociated into single cell suspension, and re-plated at 30,000 cells per well onto 24-well Transwell polyester 0.4 μm pore membrane inserts (Corning) with media filling both top and bottom chambers. Twenty-four hours after seeding, media from both chambers was replaced and 2.10^5^ freshly thawed PBMCs were added to the bottom chamber for 14 h co-culture.

CoV2 and IAV co-cultures were performed in three independent experiments (one pilot, one main experiment and one replication experiment) with at least 3 biological replicates per condition. EBOV co-culture was performed for one experiment with 3 independent replicates. Transfectant co-culture were performed in two independent experiments for each cell line (each experiment used a different preparation of plasmid DNA), and each experiment included 2 biological replicates for each transfected protein.

### Cell treatments

To determine the response to LPS, PBMCs were cultured for 14 h, in parallel to HEK co-cultures, with or without 1 ng/mL of LPS (LPS from E. coli O55:B5, Sigma, Cat# L2880) for 14 h. To assess the effects of co-culture with lysed cells, a freshly passaged single cell suspension of HEK cells at 2.5.10^5^ cells/ml was frozen on dry ice, then thawed rapidly at 37°C. The freeze-thawed suspension was centrifuged at 500 x g to eliminate debris and added at a 10-fold dilution in PBMC cultures for 14 h.

### Magnetic isolation of PBMCs in viral co-cultures

Due to lack of flow cytometry sorting in the BSL-4 containment laboratory, different PBMC subpopulations were isolated by magnetic sorting. Briefly, after 14 h of co-culture, supernatants containing PBMCs were harvested and washed, and dead cells were removed using the EasySep™ Dead Cell Removal (Annexin V) Kit (StemCell, #17899). The cell populations of interest, monocytes (CD14^+^CD16^−^) or B cells (CD19^+^), were isolated by negative magnetic selection following manufacturer’s instructions, using EasySep™ Human Monocyte Isolation Kit (StemCell, #19359) and EasySep™ Human B Cell Isolation Kit (StemCell, #17954). The isolated samples were resuspended in lysis buffer (TCL Buffer (QIAGEN) supplemented with 1% 2-mercaptoethanol) at a concentration of 500-1,500 cells per 5 µl, and frozen in a DNA Lo-Bind tube (Eppendorf, #022431021) at −80°C. For EBOV and CoV2 co-cultures, samples were heat-treated for 45 min at 60°C, removed from the BSL-4 laboratory and stored at −80°C.

In parallel, Caco-2 cells seeded in a 96-well plate were infected with one of the three viruses and cultured alone. At 48 h post infection (similar harvesting timepoint as for co-cultures), the cells were washed and directly lysed in 125 µL of TCL/βME lysis buffer, and heat-treated as above.

### Flow sorting of PBMCs in transfectant co-cultures

PBMCs were harvested after 14 h co-culture, washed twice and stained using CD14 Pacific Blue (clone M5E2, BioLegend cat# 301815, 3:100); CD19 PerCP-Cy5.5 (clone H1B19, BioLegend #302230, 3:100); CD3 Alexa Fluor 700 (clone OKT3, BioLegend, #317340, 2:100); CD45 APC-H7 (clone 2D1, BD, #560178, 2:100). Monocytes and B cells were immediately sorted as DAPI– CD45^+^CD3^−^CD19^−^CD14^+^ and DAPI-CD45+CD3-CD19+ respectively on a Moflo Astrios sorter (Beckman Coulter). 1,000 cells were sorted directly into 5 μl TCL/βME lysis buffer.

### Assessment of infection rate in epithelial cells

#### Flow cytometry

After 14 h of co-culture and the removal of the supernatant containing PBMCs, epithelial cells were harvested after trypsinisation and stained in PBS with Zombie UV viability dye (BioLegend, #423107, 1:500) for 10 min. Then, cells were fixed and inactivated in 10% formalin (Fisher Scientific) for a minimum of 6 h at 4°C and removed from the BSL-4 laboratory. Fixed cells were washed in staining buffer (PBS + 2% FBS + 1mM EDTA) and permeabilized in 0.1% saponin buffer for 10 min at room temperature (RT). After blocking the non-specific binding sites by incubation in blocking buffer (PBS + 10% of donkey serum) for 1 h, cells were stained (2 h, RT) with either: anti-GFP AlexaFluor488 (clone FM264G, BioLegend, #338008, 1:200), rabbit anti-SARS-CoV nucleocapsid (N) protein (Rockland, #200-401-A50, 1:500, cross-reacts with the CoV2 nucleocapsid protein) or goat anti-EBOV VP35 protein (custom-made by Antagene, 1:200). For EBOV and CoV2, cells were stained 30 min with the secondary antibodies anti-rabbit and anti-goat AlexaFluor488, respectively (both 1:500, Jackson ImmunoResearch). Flow cytometry was performed on a FACSymphony^TM^ flow cytometer (BD Biosciences) and analysed using FlowJo 10 software.

#### Immunohistochemistry

For CoV2 and EBOV, Caco-2 cells seeded in 96-well plates were infected as described above. One day post-infection, cells were fixed and inactivated in 10% formalin (Fisher Scientific) for a minimum of 6 h at 4°C and removed from the BSL-4 laboratory. The cells were permeabilized with acetone-methanol solution (1:1, vol:vol) for 5 min at −20°C, incubated in 0.1 M glycine for 10 min at RT and incubated in blocking reagent (2% bovine serum albumin, 0.2% Tween 20, 3% glycerin, and 0.05% sodium azide in PBS) for 20 min at RT. After each step, the cells were washed 3 times in PBS. The cells were incubated overnight at 4°C with a rabbit antibody recognizing CoV2 N protein (Rockland, #200-401-A50; 1:1000) and a goat-anti-EBOV VP35 antibody (custom-made by Antagene, 1:200) in Cov2 and EBOV infection respectively. The cells were washed 4 times in PBS and incubated with goat anti-rabbit antibody conjugated with AlexaFluor488 or donkey-anti-goat antibody conjugated with AlexaFluor594 for 1 h at RT (Invitrogen; 1:200). 4’,6-diamidino-2-phenylindole (DAPI; Sigma-Aldrich) was used at 200 ng/mL for nuclei staining. Images were acquired using a Nikon Eclipse Ti2 microscope with Photometrics Prime BSI camera and NIS Elements AR software.

For IAV, PR8-GFP-infected and PBMC-added Caco-2 cells were directly imaged live at 48 h post-infection using Nikon TE200 and SPOT imaging software (v5.2).

### Population Low-input RNA-seq

Low-input RNA-seq was performed following the standard ImmGen low-input protocol (www.immgen.org), from the 5μl of collected lysis buffer. For the viral co-cultures, magnetically isolated samples were centrifuged at maximum speed (17,000 x g) for 3 min at 4°C before loading, in order to pellet the cellular debris and magnetic particles, the cleaned RNA lysate remaining in the supernatant. Smart-seq2 libraries were prepared as described previously (4) with slight modifications. Briefly, total RNA was captured and purified on RNAClean XP beads (Beckman Coulter). Polyadenylated mRNA was then selected using an anchored oligo(dT) primer (50 – AAGCAGTGGTATCAACGCAGAGTACT30VN-30) and converted to cDNA via reverse transcription. First strand cDNA was subjected to limited PCR amplification followed by Tn5 transposon-based fragmentation using the Nextera XT DNA Library Preparation Kit (Illumina). Samples were then PCR amplified for 12 cycles using barcoded primers such that each sample carries a specific combination of eight base Illumina P5 and P7 barcodes for subsequent pooling and sequencing. Paired-end sequencing was performed on an Illumina NextSeq 500 using 2 x 38bp reads with no further trimming.

### RNA-seq data processing and QC

Reads were aligned to the human genome (GENCODE GRCh38 primary assembly and gene annotations v27) with STAR 2.5.4a (https://github.com/alexdobin/STAR/releases). The ribosomal RNA gene annotations were removed from the GTF (General Transfer Format) file. The gene-level quantification was calculated by featureCounts (http://subread.sourceforge.net/). Raw read counts tables were normalized by median of ratios method with DESeq2 package from Bioconductor (https://bioconductor.org/packages/release/bioc/html/DESeq2.html) (5) and then converted to GCT and CLS formats. To map the viral reads, the SARS-CoV2, EBOV and PR8 IAV genome sequences and annotations were obtained from NCBI, with accession numbers: GCF_009858895.2, GCF_000848505.1, GCF_000865725.1. Reads were aligned by STAR 2.5.4a, with, for SARS-CoV2, specific parameters suggested in Kim *et al*.(6).

#### Quality control

Samples with less than 1 million uniquely mapped reads were automatically excluded from normalization to mitigate the effect of poor-quality samples on normalized counts. Samples having fewer than 8,000 genes with over ten reads were also removed from the data. We screened for contamination by using known cell type specific transcripts (per ImmGen ULI RNAseq and microarray data). Finally, the RNA integrity for all samples was measured by median Transcript Integrity Number (TIN) across human housekeeping genes with RSeQC software (http://rseqc.sourceforge.net/#tin-py). For co-cultures with virally infected cells, samples with TIN <30 were removed from the BSL4 datasets prior to downstream analysis (the usual threshold, TIN <45, was used elsewhere). To eliminate unreliable datasets in the transfection analysis, samples with inter-replicate correlation <0.92 were omitted. To avoid quantitatively unreliable values from low expression, only genes with a minimum read count of 20 TPM in at least two samples were retained for analysis.

#### Differential gene expression and consensus CoV2 signature

We used an uncorrected t-test (comparing log-transformed expression values in monocytes of CoV2- vs mock-infected co-cultures) to assess differential gene expression between the different groups from the normalized read counts dataset. Genes with a FC >2 or <0.5 and nominal p-value < 0.01 were selected for some analyses (Fig. S1, Fig. S2B). A reproducible CoV2 signature set was computed by integrating the results from the first 2 co-cultures experiments (adult PBMCs with CoV2- or mock-infected Caco-2 cells). We first flagged genes which passed CoV2/Mock FC>2 and t.test −log10 (p.value) >1.5 in each experiment (and FC<0.5 for the down signatures). The intersect identified 123 transcripts matching in both experiments, but it was clear that this stringent selection, associated with the low numbers of genes reliably detected in Experiment1, were setting aside too many truly affected transcripts. We thus supplemented this list by including transcripts with CoV2/Mock FC>1.8 and correlated with the selected geneset (as mean of normalized expression across all cells) at Pearson r>0.85, yielding a cov2 signature of 429 up- and 246 downregulated transcripts, which was used as a standard in later experiments. The signature geneset was replicated to 87% in the third independent experiment. FC values (vs the mean of mock-infected control datasets) for these signature genes were imported into Morpheus (Broad Institute, https://software.broadinstitute.org/morpheus), in which k-means clustering was performed with empirically chosen k (clustering used in Fig. 2I).

#### Computation of module indices

the CoV2 index was calculated for each donor by averaging the log2 normalized expression (versus mean of all mock) of all genes belonging to the CoV2 signature (subtracting the average of the up signature to the down signature)

#### Analysis of the transfectants co-culture

Datasets from co-cultured monocytes or B cells were pre-processed as above. For Caco-2 transfectants (Fig. 3), QC-passing datasets were obtained for 27 single-protein datasets (+ GFP transfected controls), all in duplicates. We first selected transcripts with relevant variation by comparing (i) the vector of inter-replicate coefficients of variation (CV), a reliable indicator of experimental noise (computed as the mean of all inter-replicate CVs, this for every gene) with (ii) the overall CV within the dataset (averaged from CV computed from 1,000 randomly picked dataset pairs, excluding replicates). Transcripts were selected if overallCV-replicCV>0.2 OR (replicCV<0.4 & ((overallCV>(replicCV*2)), and their FC values (vs the mean of co-cultures with GFP-transfected cells) imported into Morpheus for heatmap representation and clustering (Fig. 3B, split into 7 empirically determined k-means clusters). The same procedure was followed for HEK transfectants (Fig. S3B).

#### Correlation analysis of transfected Caco-2 cells with corresponding co-culture effects

To search for transcripts in transfected Caco-2 cells associated with induction of the CoV2-Up signature in co-cultured monocytes, we first constructed a coculture effect vector by computing the mean log2 fold change vs GFP controls of genes belonging to the CoV2-Up signature in monocytes co-cultured with Caco-2 cells transfected with individual viral genes. Using matched transfectant samples, we computed a per-gene correlation (Spearman) between this vector and the log2 fold change in Caco-2 cells transfected with individual viral genes vs GFP controls. To assess the statistical significance of the resulting correlation coefficients, we repeated this procedure after permuting sample labels 100 times. As the correlation coefficients from the permutation were approximately normally distributed, p values were calculated using a two-tailed one-sample z-test using the mean and standard deviation of the coefficients from permuted data for each gene.

#### Geneset enrichment analysis

Enrichment of CoV2.up and CoV2.dn signature in Gene Ontology (GO) biological processes pathways and Reactome pathways was computed using Fisher’s exact test with BH-FDR correction, through the g:Profiler interface. Pathways with a false discovery rate (FDR) <1% were selected for further analyses and recorded in Dataset S2. In order to overcome the inherent redundancy of GO pathways, significant pathways were then interpreted and visualized as an enrichment network using Cytoscape (v3.8.2) (7) and its *EnrichmentMap* and *AutoAnnotate* modules. Briefly, Cytoscape allows collapsing of redundant significant pathways into a single biological theme using the Jaccard Overlap combined index (cutoff=0.5), thus simplifying the interpretation. After filtering noise, the network represents overlaps among the most enriched pathways (FDR 0.5%), in which similar pathways are automatically group into main biological themes. Regarding the type I IFN and IFN gamma signatures, data were downloaded from the Gene Expression Omnibus (GEO) (https://www.ncbi.nlm.nih.gov/geo/) from two human published datasets GSE142672 and GSE46599 (8). To reduce noise, genes with a CV between biological replicates <0.7 and an expression level >20 in either comparison groups were selected. Upregulated transcripts were defined, at an arbitrary threshold, as having a FC >1.5 and a t-test p-value <0.05.

#### Overlap with COVID-19 patient profiling datasets

Signatures from *in vivo* myeloid population were extracted from published sources. The alveolar macrophage signature from severe COVID-19 patients was extracted from Liao et al. (9), by merging the signature of their two predominant populations found in severe patients: FCN1^hi^ (group1) and FCN1^lo^SPP1^+^ (group2). Signature of the MS1 state in sepsis was directly downloaded from the supplementary tables of Reyes et al (10). PBMC datasets from (11) were retrieved from the COVID19 cell atlas at https://www.covid19cellatlas.org/index.patient.html as an R dataset object which included the cellXgene matrix, and were used with the cell annotation provided by Wilk et al. The CD14^+^ monocyte cluster was extracted; indices for the CoV2-up signature genes were computed by averaging the pre-computed CoV2-up genes together per cell, and color-coded on the monocyte UMAP space for Fig.4C.

### Statistical analysis

Unless specified otherwise, the data are presented as mean ± SD and tests of associations for different variables between infected/transfected and mock/GFP were computed using the nonparametric Mann Whitney test. Significance of signature overlaps into our dataset was assessed by Chi square test when computing one signature at a time (e.g. assessing one signature in an independent volcano plot), or by a Fisher’s exact test with BH-FDR correction when using large curated GO signatures databases. Analyses and plots were done using S-Plus (v8.2.0), RStudio (v.1.2.5019) and GraphPad Prism (v.8.4.3), heatmaps generated with Morpheus (https://software.broadinstitute.org/morpheus).

### Data availability

The data reported in this paper have been deposited in the Gene Expression Omnibus (GEO) database under accession no. GSEXXX (human RNAseq).

## SUPPLEMENTARY FIGURE LEGENDS

**Fig S1.**
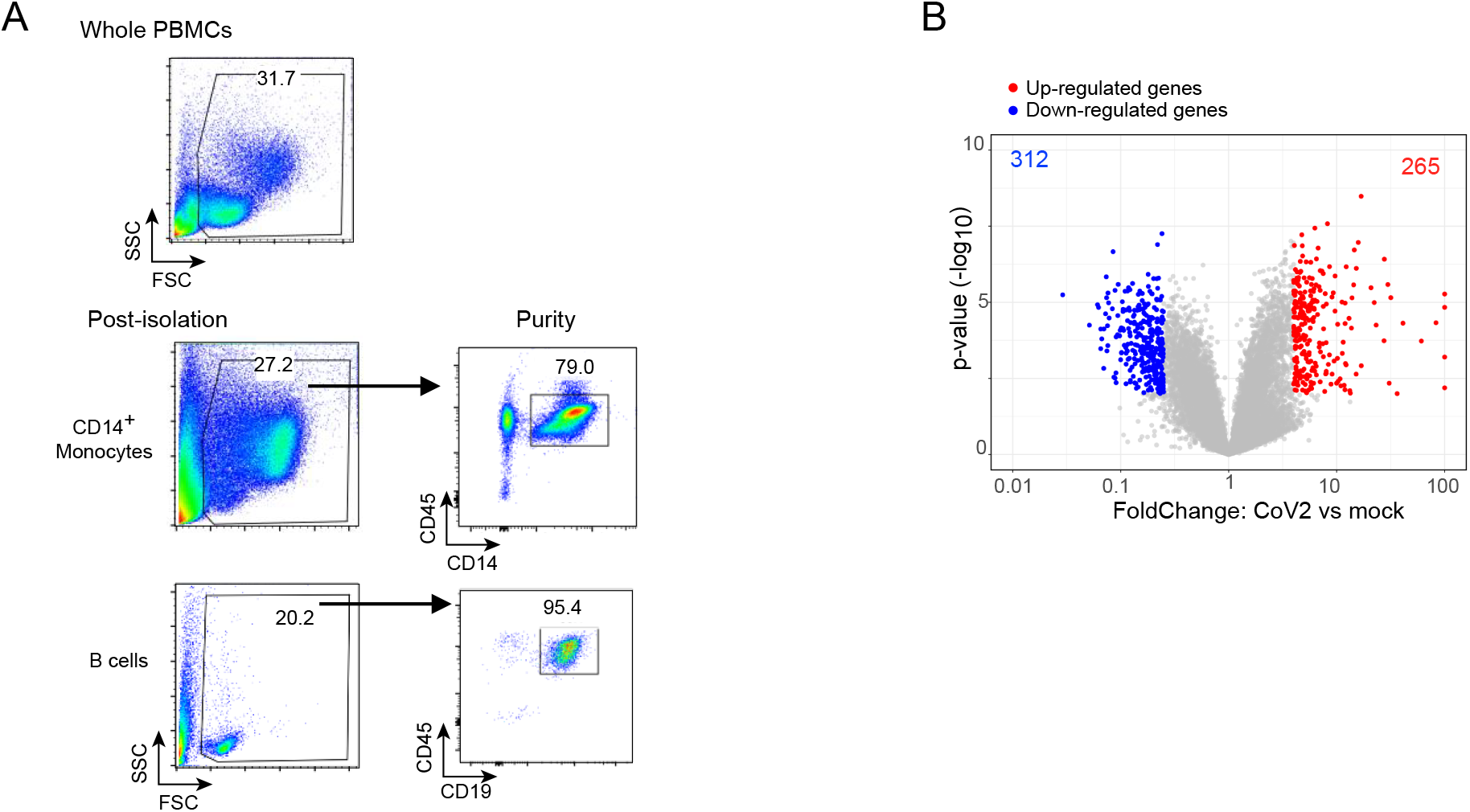
**(A)** Representative flow cytometry plots showing a sample of the final isolated fractions of magnetically-purified monocytes (top) and B cells (bottom). Data from one set-up co-culture experiment. **(B)** Volcano plots displaying gene expression changes in CoV2-infected Caco-2 cells versus mock Caco-2 cells, 48 h post-infection. On the left panel are highlighted the differentially expressed genes (p<0.01, FC>2 or <0.5). On the right panel are highlighted the CoV2 upregulated (red) and downregulated (blue) gene signatures. P-values were calculated using the Chi square test. **(C)** Volcano plot of gene expression in B cells co-cultured with CoV2 infected Caco-2 compared to monocytes co-cultured with uninfected Caco-2 (mock). Differentially expressed genes are highlighted (p<0.01, FC>2 or <0.5).

**Fig S2.**
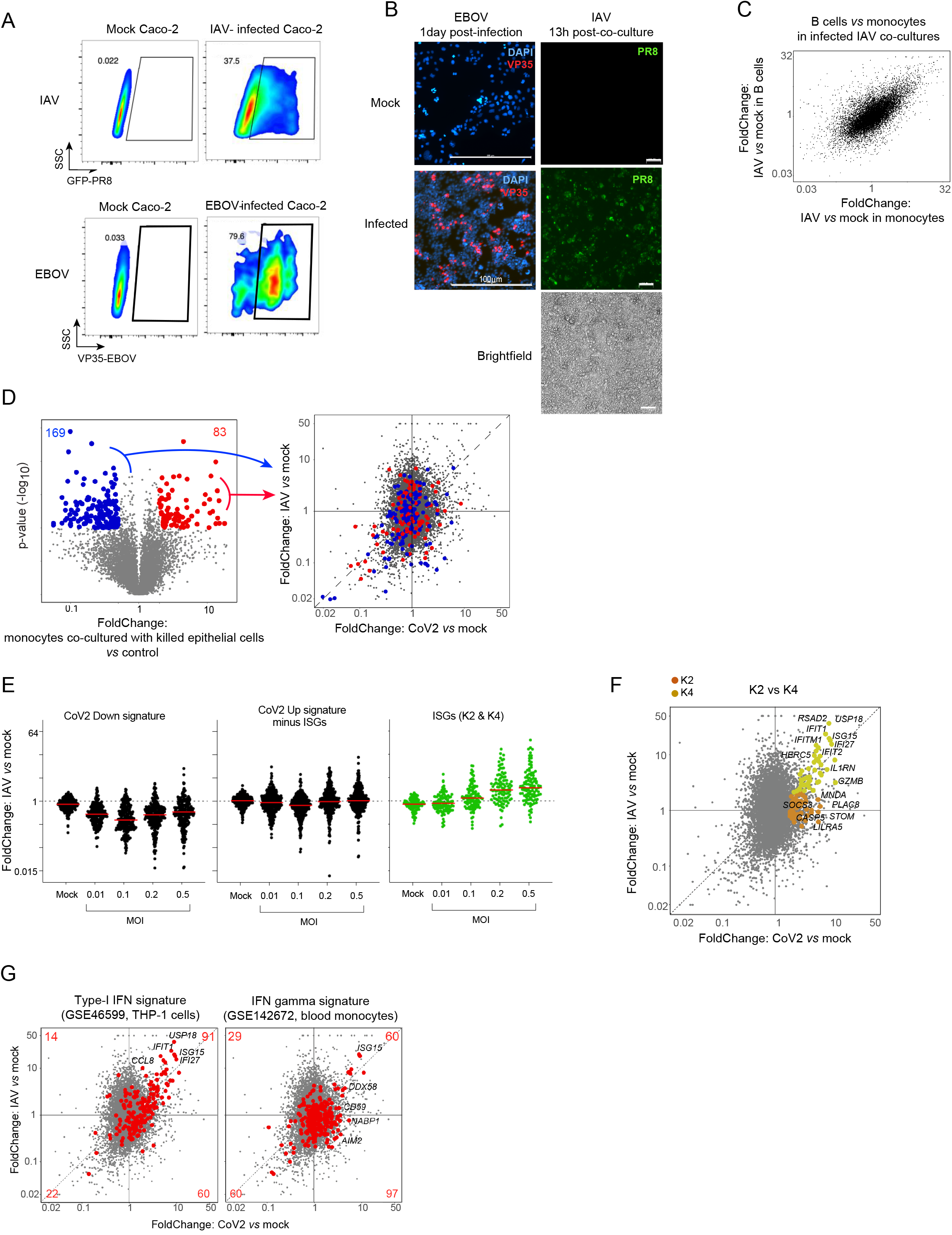
(**A-B**) Infection rate of IAV and EBOV in Caco-2 cells assessed: **(A)** by flow cytometry at the time of harvesting PBMCs (∼48 h post-infection, one sample of the third independent experiment for IAV) **(B)** by immunofluorescence microscopy at 1 day post-infection for EBOV (left panel, 10X magnification) or at the time of harvesting PBMCs for IAV (∼48 h post-infection in the setting of co-culture PBMCs added to infected epithelial cells, one sample of the second independent experiment, right panel). Infected cells are marked in green (GFP-PR8) for IAV and in red (anti-VP35 + anti-goat-AF594) for EBOV. Nuclei are marked with DAPI (blue, EBOV) or cellular membranes visible in brightfield (IAV). (C) FoldChange-FoldChange plot comparing the response of B cells versus monocytes in the context of IAV infection in Caco-2. (D) Overlay of transcriptional signature derived from monocytes co-cultured with killed epithelial cells onto viral co-culture datasets. Left panel shows the gene expression changes induced in monocytes co-cultured with killed epithelial cells (freeze-thawed HEK, see methods) highlighting the differentially expressed genes (p<0.01, FC>2 or <0.5). Right panel shows these genes into the FoldChange-FoldChange plot CoV2 versus IAV in monocytes. (E) Changes in gene expression as a function of MOI of IAV infection in monocytes co-cultured with IAV infected Caco-2. Each gene is a dot. Genes belonging to the CoV2 down signature (left panel), the CoV2 up signature excluding the 2 ISG clusters (middle panel) or the two ISGs clusters (right panel) are shown. (F) Dichotomy in the IFN response between IAV and CoV2 highlighted in a FoldChange-FoldChange plot comparing the response in monocytes co-cultured with IAV (y-axis) relative to CoV2 (x-axis) infected Caco-2. K2 (orange) and K4 (yellow) are highlighted. (G) Same plots than in (F), highlighting: upregulated transcripts from GSE46599 (type I IFN-treated THP-1 cells for 24h), left panel, and GSE142672 (IFN gamma-treated blood monocytes for 24h), right panel.

**Fig S3.**
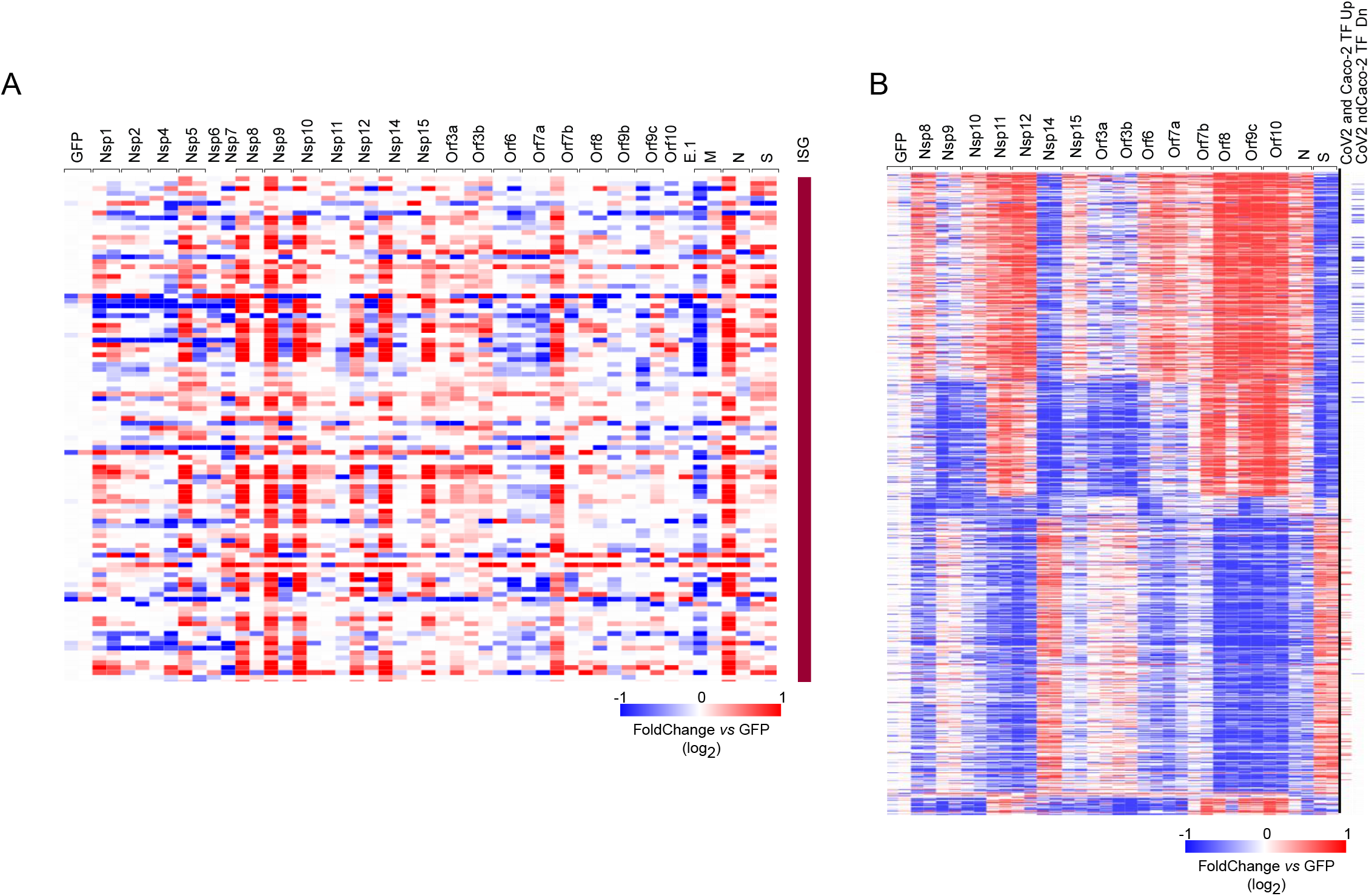
(**A**) Heatmap of ISG transcript expression values in the Caco-2 transfectant dataset. (**B**) Heatmap of the significantly differential expressed transcripts in monocytes co-cultured with transfected HEK (similar selection as for Caco-2: overall variance in the dataset substantially exceeded inter-replicate variance, and with significant difference from GFP-transfected controls in at least one co-culture FC>2 or <0.5, p<0.01). The columns represent the different conditions, where only proteins for which both replicates passed the QC are shown. Annotations at the right ribbon show the overlap between these genes and part of the CoV2 signature that was up/down regulated in Caco-2.

**Fig S4.**
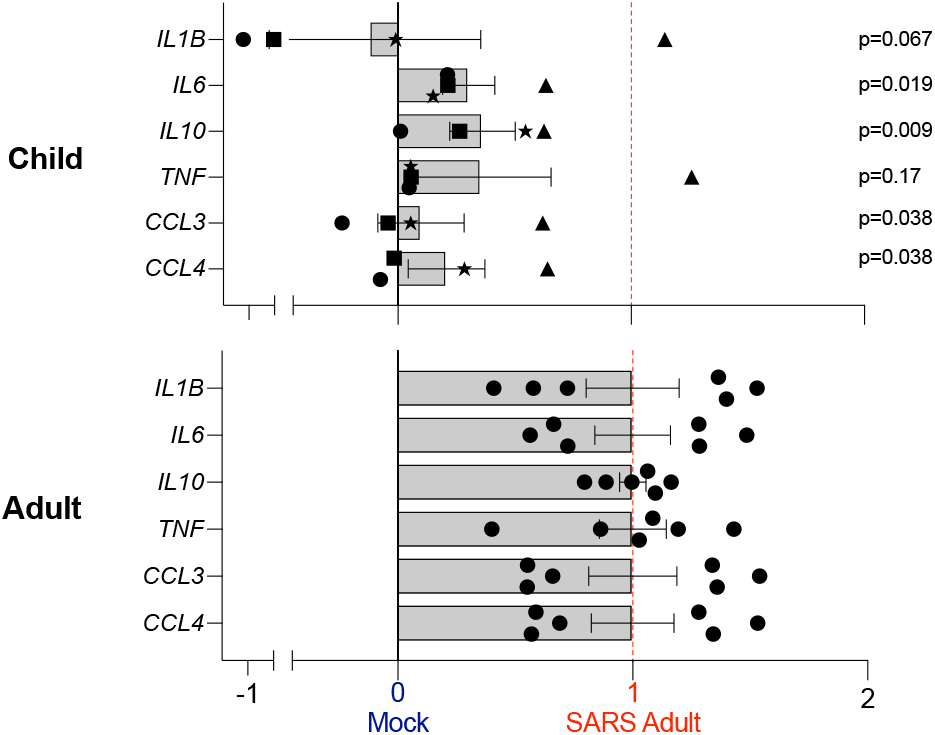
Representation of the average expression of a selected set of cytokines across all CoV2-co-cultured monocyte replicates (mean and SEM), child (upper panel) or adult (lower panel). Each individual replicate is a dot. Score computed independently for each cytokine, where 0 corresponds to the average expression of this cytokine in the respective mock condition and 1 its average expression in CoV2-co-cultured adult monocytes (red line). The annotated p-values were computed from the values in children vs adult samples for each cytokine, using the Mann–Whitney test.

## SUPPLEMENTARY TABLE LEGENDS

**Table S1**: Cov2 signature

**Table S2**: Gene ontology analyses of the CoV2 signature

**Table S3**: Donor information and viral reads among the different samples and conditions

**Table S4**: Heatmap of the CoV2 signature among all CD14+ samples (Fig. 2I - FoldChange vs the average of the respective mocks)

